# Neural mechanisms underlying distractor suppression guided by spatial cues

**DOI:** 10.1101/2022.05.22.492997

**Authors:** Chenguang Zhao, Yuanjun Kong, Dongwei Li, Jing Huang, Xiaoli Li, Ole Jensen, Yan Song

## Abstract

A growing body of research demonstrates that distracting inputs can be proactively suppressed via spatial cues, nonspatial cues, or experience, which are governed by more than one top-down mechanism of attention. However, how the neural mechanisms underlying spatial distractor cues guide proactive suppression of distracting inputs remains unresolved. Here, we recorded electroencephalography signals from 110 subjects in three experiments to identify the role of alpha activity in proactive distractor suppression induced by spatial cues and its influence on subsequent distractor inhibition. Behaviorally, we found novel spatial changes in spatial distractor cues: cueing distractors far away from the target improves search performance for the target while cueing distractors close to the target hampers performance. Crucially, we found dynamic characteristics of spatial representation for distractor suppression during anticipation. This result was further verified by alpha power increased relatively contralateral to the cued distractor. At both the between- and within-subjects levels, we found that these activities further predicted the decrement of subsequent P_D_ component, which was indicative of reduced distractor interference. Moreover, anticipatory alpha activity and its link with subsequent P_D_ component were specific to the high predictive validity of distractor cue. Together, these results provide evidence for the existence of proactive suppression mechanisms of spatial distractors, support the role of alpha activity as gating by proactive suppression and reveal the underlying neural mechanisms by which cueing the spatial distractor may contribute to reduced distractor interference. (235).

**Significance:** In space, the attention-capturing distractors are obstacles to successfully identifying targets. How to sidestep task-irrelevant distractors that stand between the target and our focus in advance is essential but still unclear. This research investigated how dynamic spatial cues can help us proactively eliminate attention-capturing distractors. Using three cue-distractor tasks that manipulate the predictive validity of distractor occurrence, we provide a series of evidence for the presence of alpha power activity related to distractor anticipation. Critically, this was the first study linking cue-elicited alpha power and distractor-elicited PD, indicating that spatial modulation of alpha power may reduce distractor interference. These findings delineate the neural mechanisms of proactive suppression for spatial distractors. (109)

## Introduction

In daily life, individuals often select a task-relevant target from the surrounding distractors, for which selective attention is required (Luck and Hillyard, 1994). Competition of simultaneously presented distractors for limited attentional resources is likely to be inhibited in advance via a “proactive suppression” mechanism (Geng et al., 2014). The emerging consensus on the mechanism of proactive suppression is flexible and not unitary (van Moorselaar et al., 2020; Noonan et al., 2016), it might be influenced via contextual factors (Marini et al., 2013; 2016), statistical learning (Wang et al., 2018; 2019), and nonspatial features (Snyder et al., 2010; Gutteling et al., 2022). However, the jury is still out on how spatial distracting information can be filtered out proactively.

Prior behavioral work provided mixed evidence for proactive suppression. Several studies have shown that giving the distractor-related location in advance is likely to harm (Moher et al., 2012; Tsal et al., 2006) or benefit the target response (Munneke et al., 2008; Chao et al., 2010). Some behavioral research shows that proactive suppression might not take place unless the location of the upcoming distractor becomes predictable by repeating stimuli or blocked design (Cunningham et al., 2016; Wang et al., 2018). However, recent research suggests that distractors can be suppressed proactively in trialwise analysis by showing that eye movements are less likely to be deployed to a cued distractor (van Zoest et al., 2021).

Given the tight link between spatial attention and alpha power, spatial attention bias is usually measured by alpha-band power activity (8-12 Hz). A substantial body of work has linked alpha-band activity to spatial suppression, and recent studies have mainly focused on two guises of alpha power originating from distinct research traditions: hemispheric lateralization and spatial selectivity as will be reviewed below.

First, cue-elicited hemispheric lateralization of alpha power over posterior cortices is considered a signature of active attention control (Thut et al., 2006). Alpha power relatively increased contralateral to to-be-suppressed irrelevant visual inputs termed the “negative” alpha modulation of distractors (Zhao et al., 2022). Importantly, such alpha power lateralization was observed before the distractor onset on a trial-by-trial basis, which speaks to alpha lateralization as proactive suppression for upcoming distracting inputs (Wöstmann et al., 2019). Such alpha lateralization reflects the distractor-related bias of spatial attention, which is interpreted as gating by the distractor inhibition hypothesis (Jensen and Mazaheri, 2010).

Second, an influential line of research focused on the spatial selectivity presented by alpha activity. An inverted encoding model (IEM) was applied to track the temporal and spatial dynamics of spatial attention (Foster et al., 2017 a,b; Samaha et al., 2016; Popov et al., 2019). Spatial distribution of alpha power across electrodes, termed a channel tuning function (CTF), enabled a more refined selectivity of the attention bias. Indeed, the fact that the spatially distributed alpha activity precisely tracks the position of the target, even in the absence of irrelevant distractors, casts doubt on whether the functional role of alpha oscillations is consistent with the distractor inhibition hypothesis (Foster et al., 2019). Several studies suggest that the evidence for alpha power as a distractor inhibition account is limited (Foster et al., 2019), thus it is debated to what extent alpha oscillation can proactively suppress distractors (Noonan et al. 2018; van Moorselaar & Slagter, 2020; van Zoest et al. 2021).

Beyond alpha activity, distractors can elicit one ERP component, P_D_ (positive distractor), which has been proposed to reflect reactive prevention or termination of salient distractors. The decreased amplitude of the P_D_ is thought to reduce distractor interference in spatial priority maps (Gaspar et al., 2014). To date, P_D_ amplitude can be influenced by learned suppression (van Moorselaar et al., 2019; 2020), nonspatial suppression (Arita et al., 2012), and strategy (van Zoest et al., 2021). There is still a lack of evidence for spatial suppression.

To address these unsettled issues, we manipulated a variant of the Posner paradigm (Fig. 1A) by using spatial circular radar-like cues, where given prior spatial information was informative or uninformative (Experiment 1), with further manipulation for the validity of information (Experiment 2), and symbolic alternation (Experiment 3). Through three EEG experiments, we aimed to investigate cue-induced alpha activity and the P_D_ elicited by distractors, as well as their link. We assume that if proactive suppression of the upcoming distractor is related to alpha activity, corresponding changes in alpha power should be observed following different spatial cues, and it can explain the variance in P_D_ amplitude. (738).

**Fig. 1.**
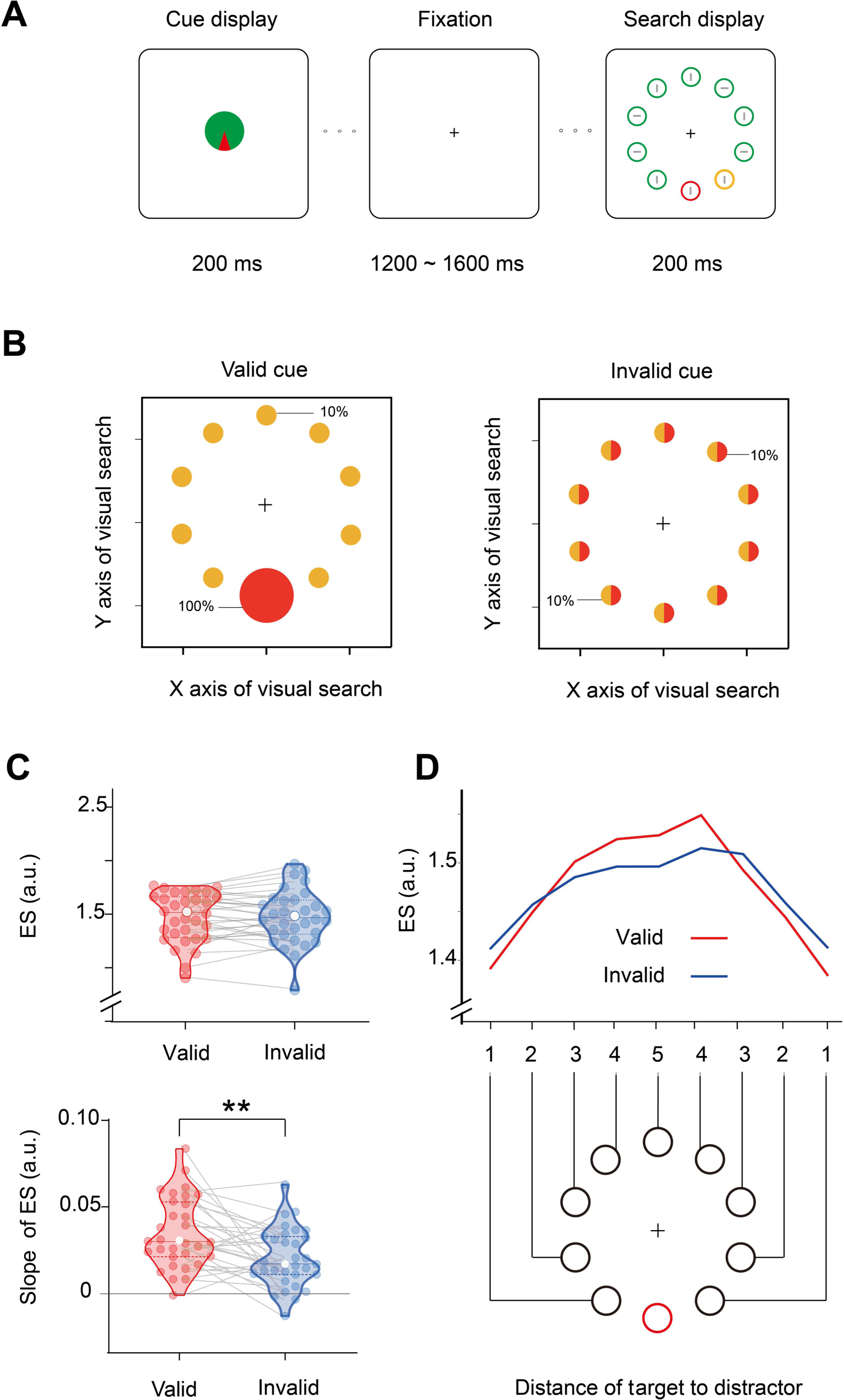
Task paradigm and behavioral results for Experiment 1. **A** Each trial began with a cue display of the distractor, 1200∼1600 ms followed by a search display. In two separate sessions, the cue display was fully predictive (with 100% validity) or not predictive (with 10% validity) of the specific location of the red distractor circle. Participants were instructed to indicate the orientation of the gray line inside the yellow target circle in the search array. **B** The spatial probability of the target and distractor occurring during subsequent visual search with respect to two cue sessions (yellow represents the target; red represents the distractor). **C** The mean (top) and slope (bottom) of ES in the valid (red) and invalid (blue) cued distractor sessions. Violin plots depict the distributions of measurements in each session, with dots representing each subject. The solid and dotted lines indicate medians and quartiles, respectively. **p < 0.01 **D** The diagram illustrates the changes in ES across the distances of the target to the distractor location (DTD). The red circle indicates the distractor, and the black circles indicate potential targets at different distances to a distractor.

## Results

The materials and methods of these experiments are available at the Open Science Framework (https://osf.io/z9rym/).

### Experiment 1

To study the neural mechanisms underlying distractor suppression guided by spatial cues, we performed two sessions in Experiment 1 (see Fig. 1A). For the valid-cue session, the radar-like cue was fully predictive of the direction in which the subsequent distractor would appear (red represents the distractor; yellow represents the target). We also included an invalid-cue session in which the distractor location was uninformative for invalid-cue sessions. The comparison between invalid-cue session and valid-cue session allowed us to assess spatial attentional bias related to cued direction.

#### Behavior

To control for speed-accuracy trade-offs, we used efficiency scores (ES) by dividing the accuracy rate by reaction time (larger scores mean more efficient responses). The mean ES for the two sessions in Experiment 1 is shown in Fig. 1C (top panel). No significant distractor cueing effects (valid − invalid) of mean ES were found (t_29_ = 0.087, p = 0.931, BF_10_ = 0.190, two-tailed, Cohen’s d = 0.015). To examine spatial changes of behavior outcomes, the trial was divided into nine subgroups according to the relative distances of the target to the distractor location (abbreviated as DTD). The catalog is listed clockwise around the imaginary ring in Fig. 1D. Then, the ES of each subgroup was averaged to examine the response to the target when the distractor appeared at different DTDs (Fig. 1D). Repeated measures ANOVA on mean ESs showed a significant main effect of DTD for the valid-cue sessions (F8, 232 = 32.56, p < 0.001, η^2^ = 0.512) and the invalid-cue sessions (F8, 232 = 16.44, p < 0.001, η^2^ = 0.347).

To quantify the extent of the ES scales changed with DTD, the slope was characterized by collapsing trials across the same DTD and fitting these data by a linear function (Default function of MATLAB: polyfit.m). The slope was significantly larger than zero for the valid-cue session (t_29_ = 9.699, p < 0.001, two-tailed, Cohen’s d = 1.715) as well as for the invalid-cue session (t_29_ = 6.799, p < 0.001, two-tailed, Cohen’s d = 1.202). These results show a spatial distractor interference in which the salient distractor might interfere more with target processing when the distractor singleton was present closer to the target location and vice versa, which is consistent with a previous report (Wang et al., 2018).

Interestingly, we found a significant distractor cueing effect of the slope of ES (t_29_ = 3.356, p = 0.002, two-tailed, Cohen’s d = 0.593) in Experiment 1, as expected (see Fig. 1C, bottom panel). Compared with the invalid-cue session, we found that participants had better performance when distractors occurred at locations (the 5^th^ and 4^th^) far away from the target in the valid-cue session. In contrast, participants had poorer performance when distractors occurred at locations (1^st^ and 2^nd^) near the target. If the distractor cueing effect has a spatial extent, we expect that the slope of ES in the valid-cue session may be steeper than that for the invalid-cue session. We obtained a similar cueing effect in participants’ reaction time and accuracy (Fig. S1): A significant cueing effect was found on the slope of accuracy (t_29_ = 2.556, p = 0.016, two-tailed, Cohen’s d = 0.452) and reaction time (t_29_ = −2.579, p = 0.015, two-tailed, Cohen’s d = −0.456). Taken together, our preliminary results showed novel spatial behavioral changes, which supported the existence of proactive suppression for spatial distractor cues.

#### Alpha channel-tuning function (CTF) of distractor cueing

Previous research suggested that spatial distribution of neural representation was especially pronounced within the alpha band power (8 to 12 Hz). The inverted encoding model (IEM) analysis was applied to reconstruct the attended distractor location from the pattern of alpha power to explore spatial selectivity. As shown in Fig. 2A, this procedure produces CTFs, which reflect the spatial distribution of alpha power that is measured by scalp EEG (conceptualized into ten ideal electrodes). In brief, the center channel was tuned for the position of the direction of interest (e.g. 180° red arrow in Fig 2A left), then channel offsets (e.g. 72° in Fig 2A right) were defined as the angular difference between the center channel and other channels. Each estimated CTF was then circularly shifted to a common center (0° on the channel offset axes of Figure 2B) and several channel offsets (−180° to 180°). The final CTF was a function associated with the shifted channel offsets.

**Fig. 2.**
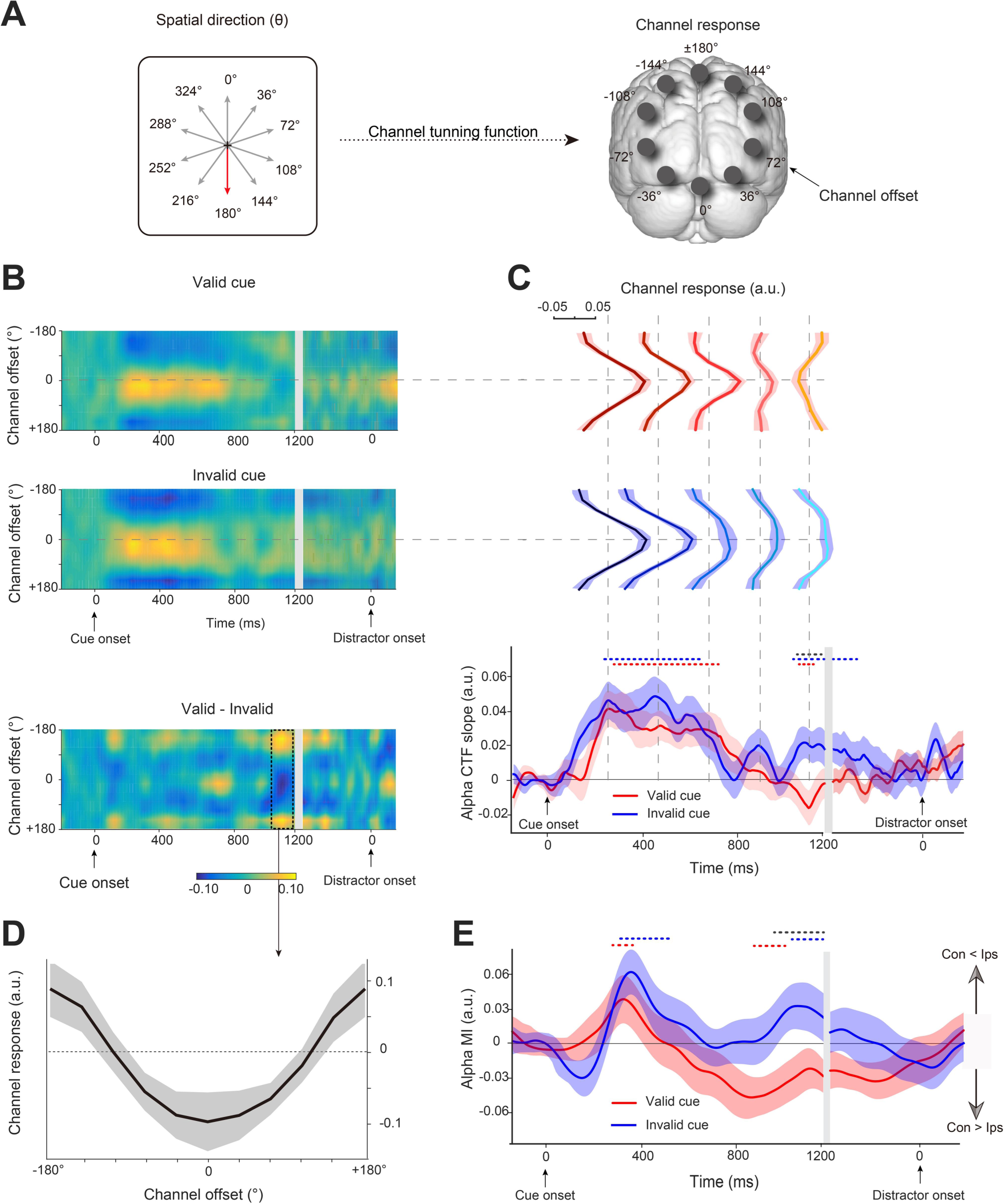
EEG results during the cue-distractor intervals from Experiment 1. A The spatial direction of the distractor cue varied from trial to triaL The spatial distribution of alpha power was modeled by the channel tuning functions across ten ideal channel offsets, right panel show channel offsets and the centre channel if distractor cue point 180 degrees (red arrow). **B** Alpha-band CTFs across the cue-distractor intervals for valid-cue and invalid-cue sessions. The difference between the two sessions was also plotted. C The direction selectivity of the alpha-band CTF (measured as CTF slope) across time in valid (red) and invalid (blue) cue sessions. The different channel response curves at five sampled time points (gray vertical dashed lines) were plotted in both sessions. D The cueing effect on alpha-band CTFs (valid-invalid; averaged from 1040 to 1200 ms) is related to anticipation of the distractor. E Time course of the alpha MI in the posterior electrodes for valid (red) and invalid (blue) cue sessions. The red and blue dashed lines indicate a significant difference from 0, and the black dashed line indicates clusters with a significant difference between two sessions (p < 0.05). Shades of light color along with the dark color lines represent error bars (±1 SEM). Con: contralateral to distractor cue; Ips: ipsilateral to distractor cue.

Fig. 2B shows CTFs across the cue-distractor intervals (cue-locked: −200 to 1200 ms; distractor-locked: −400 to 200 ms) for valid- and invalid-cue sessions. To measure the spatial selectivity of channel responses, the time-resolved slope of CTFs was calculated for both the valid (red lines in Fig. 2C) and invalid cue sessions (blue lines in Fig. 2C). Channel response curves were plotted at different sampled time points from the maximum (T1: 224 ms) to the minimum (T5: 1136 ms) of the slope of CTFs for the valid-cue session and a set of equal diversion points between T1 and T5 (T2: 452 ms, T3: 680 ms, T4: 908 ms). These results suggest that CTFs are sensitive to the attended distractor location and time course, which was tracked by the spatial response of alpha power across different channel offsets.

In both valid- and invalid-cue sessions, the distractor selectivity (positive slope of CTFs) shows an initial rapid rise in tension followed by a gradual decrease, resulting in a slope significantly (permutation test: p < 0.050, two-tailed) different from zero in valid-cue sessions (from 264 to 744 ms) and invalid-cue sessions (from 196 to 648 ms). This shows that the alpha power was first selective for cued direction regardless of whether it had distractor-related information. Then, invalid cues still led to a significant slope from 1064 to 1200 ms locked to cue display and from −400 to −236 ms locked to the distractor (permutation test: p < 0.050, two-tailed), suggesting that channels continued to be selective for the cued location in invalid-cue sessions.

Given that the positive slope of CTFs represents the selectivity of neural activity responses to cued distractor location, the negative slope may represent the suppression of neural activity responses to the distractor. In contrast, valid cues led to distractor suppression (negative slope of CTFs) from 1062-1168 ms (Fig. 2C, bottom panel; permutation test: p < 0.050, two-tailed), resulting in a significant distractor cueing effect on the slope of CTFs from 1040 to 1200 ms between the valid-cue session and the invalid-cue session (permutation test: p < 0.050, two-tailed). The mean difference in CTFs (valid − invalid) in the significant time windows was averaged to identify the change in the channel response curve. As shown in Fig. 2D, channel response relatively increased at electrodes far away from cued distractor location, channel response relatively decreased at electrodes close to cued distractor location.

Together, our results show dynamic spatial alpha power tuning to the cued distractor location during the cue-distractor interval. The negative CTF slope was only observed in valid-cue sessions, which indicates that cueing distractors might suppress spatially subsequent distracting input by flipping the spatial tuning to the attended distractor location in advance.

#### Alpha MI of distractor cueing

Then, our interest lay in specific spatial distribution effects of alpha power—lateralized alpha power. This lateralized alpha power is defined as the difference between the alpha power in the contralateral hemisphere and that in the ipsilateral hemisphere with respect to distractor and is usually measured by the alpha modulation index (MI; Vollebregt et al., 2015; Zumer et al., 2014). To enable isolation of lateralized distractor-specific alpha power, the alpha MI evoked by cues was computed based on trials where the cue point was four of ten possible directions (288° 108 ° 252 ° 72 °). Trials were categorized as left-cued when they pointed 288° or 252°, whereas those that pointed 108° or 72° were classified as right-cued trials. Then, we combined alpha band power for left-cued trials minus right-cued trials, normalized by their mean, and averaged over left and right (see Materials and Methods for details).

As shown in the time-course representation in Fig. 2E, mimicking the CTF findings, our results show that the amplitude of MI was significantly positively modulated during the 244–560 ms and 1012–1200 ms invalid-cue sessions (permutation test: p < 0.050, two-tailed). The alpha MI in valid-cue sessions showed a significant positive modulation from 208 to 348 ms and a significant negative modulation during the late period from 748–1012 ms (permutation test: p < 0.050, two-tailed). Testing for distractor cueing effects revealed a significant difference between valid- and invalid-cue sessions during the late period of 888–1200 ms (p < 0.050, two-tailed). This result suggested that for the valid cue, the alpha power was more strongly elevated over the hemisphere contralateral to the cued distractor field during later

#### Distractor-elicited PD

We then focused on the ERPs during the subsequent visual search display. We only used trials with a lateral distractor and midline target present, in which a lateral distractor can evoke P_D_ components. The P_D_ component was present as a positive deflection in the ERP waveform at the visual cortex contralateral relative to ipsilateral to the distractor. The shaded area in Fig. 3A shows difference waveforms (contralateral > ipsilateral), revealing that a significant P_D_ (248−316 ms; permutation test: p < 0.050) was apparent at P7/8 electrodes in the invalid-cue sessions (t_29_ = 2.228, p = 0.034, two-tailed, Cohen’s d = 0.414) but not in the valid-cue sessions (t_29_ = −0.007, p = 0.995, two-tailed, Cohen’s d = −0.001), which resulted in a significant P_D_ difference between the two sessions (t_29_ = −2.090, p = 0.046, two-tailed, Cohen’s d = −0.388). These results were consistent with prior work reporting distractor learning-related reductions in P_D_ amplitude (van Moorselaar et al., 2019). Our results suggested that cueing distractors appeared to be a reduced need to reactively inhibit the capture of salience distractors, as evidenced by reduced P_D_ amplitude.

**Fig. 3.**
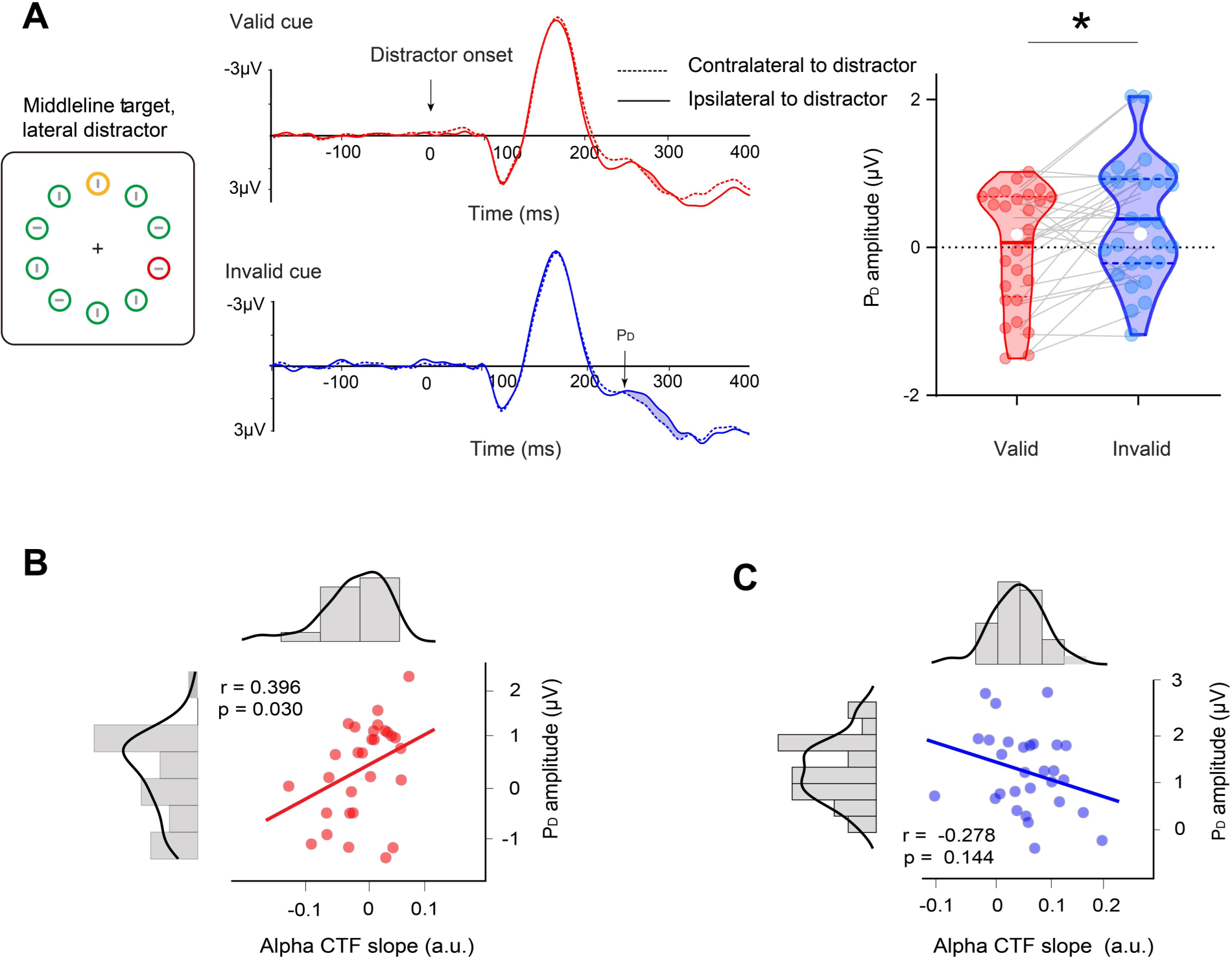
A ERP results during the stimulus period from Experiment 1. Grand averaged ERPs at contralateral and ipsilateral electrode sites relative to the distractor (averaged over P7 and P8) in valid- (red) and invalid- (blue) cue sessions. Violin plots depict the P_D_ amplitude (248-316 ms) in the two sessions, with the dots representing each subject. The solid and dotted lines indicate medians and quartiles, respectively. *p < 0.05. **B** Alpha CTF slope during the cue period as a function of the subsequent distractor-elicited P_D_ amplitudes during a visual search between participants in the valid session. The diagrams along with the scatter plot are the frequency distributions of alpha CTF slope and P_D_ amplitude, respectively. **C** Scatter plot for invalid sessions. Con: contralateral to distractor cue; Ips: ipsilateral to distractor cue.

We also conducted decoding analyses with ERP waveforms across all electrodes. The decoding performance for distractor locations did not show a difference between valid- and invalid-cue sessions (Fig. S2, left panel, permutation test: p > 0.050). We also compared decoding performance between the target and distractor (see Fig. S2). The maximum value of decoding performance for distractors was much smaller than the target decoding performance even based on the same EEG data (permutation test: p < 0.001, two-tailed), which indicated that the weight of activities related to distractor suppression was much smaller than that related to target processing.

We used between-subject correlation analysis to investigate the relationship between cue-induced alpha activity and subsequent distractor-elicited P_D_ in valid- and invalid-cue sessions. We first correlated the slope of alpha CTF (1064 to 1188 ms) with the PD. A significant correlation was found for the valid-cue sessions (r = 0.396, p = 0.030, Fig. 3B), but no significant correlation was found for the invalid-cue sessions (r = 0.278, p = 0.144, Fig. 3C). These results showed that negative alpha CTF was related to reduced distractor-elicited PD. We also correlated alpha MI with the PD, and no significant correlation was observed for the valid-cue session (r = 0.311, p = 0.454) or invalid-cue session (r = 0.151, p = 0.576). This result may be due to the relatively small number of trials performed in alpha MI analysis, of which part have no adequate power to explain the variance of whole sample PD. Behaviorally, we further investigated whether the slope of ES is correlated with the slope of CTF or the amplitude of P_D_ in Experiment 1, no correlation was found in different sessions (ps > 0.684).

### Experiment 2

In Experiment 2 (see Fig. 4A), cue pointed left or right, and informed the participants of the approximate scope in which the upcoming distractor would occur in the search display, instead of the exact location in Experiment 1. The variable scope of the distractor cue across three trials was related to the predictive validity of distractor occurrence. This manipulation of predictive validity allowed us to exclude the possibility that the hypothesized evidence for proactive suppression in Experiment 1 simply reflects the information gap between the informative cue (valid) and uninformative cue (invalid).

**Fig. 4.**
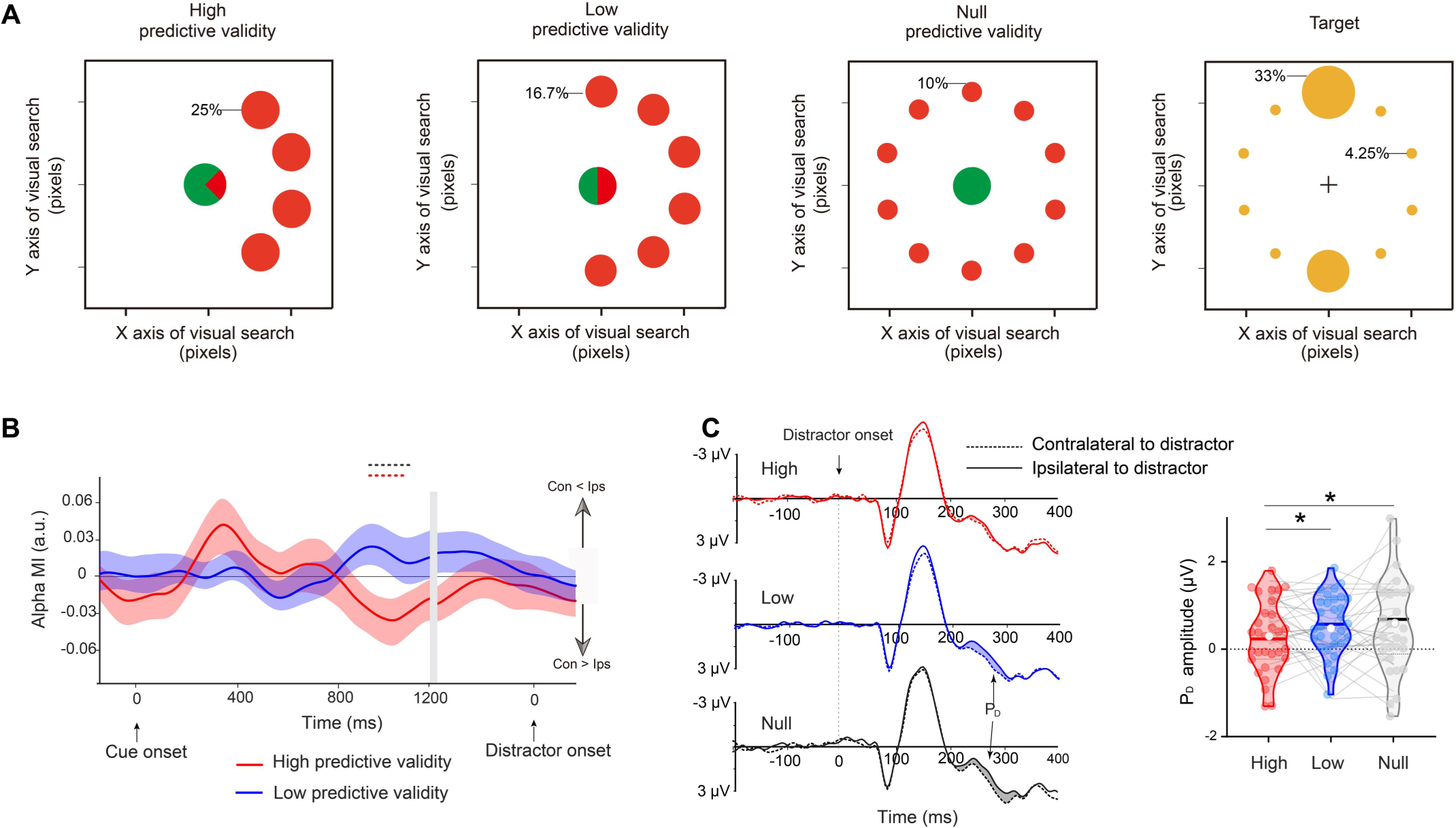
Task paradigm and EEG results for Experiment 2. **A** Three types of cue displays and corresponding spatial probability of the target and distractor occurring during subsequent visual search. Note that spatial probability was conceptual and did not actually appear around the cue. **B** Time course of the alpha MI in the posterior electrodes for high- (red) and low- (blue) predictive validity trials. Shades of light color along with the dark color lines represent error bars (±1 SEM). **C** Grand averaged ERPs at contralateral and ipsilateral electrode sites relative to the distractor (averaged over P7 and P8) in high- (red) and low- (blue) predictive validity sessions. Violin plots depict the P_D_ amplitude (248-316 ms) in the two sessions, with the dots representing each subject. *p < 0.05. Con: contralateral to distractor cue; Ips: ipsilateral to distractor cue.

Experiment 2 adopted the methods and indicators of Experiment 1 for behavioural analysis. Note that spatial probabilities of the target were not uniform across spatial location but remained equivalent among the three cue trials (see Fig. 4A), which allowed us to compare responses for the target with variable spatial probabilities of a distractor. As in Experiment 1, behavioral results (Fig. S3) showed a main effect of predictive validity on mean ES (F2, 50 = 6.449, p = 0.003, η^2^ = 0.205) and slope of ES (F2, 50 = 34.480, p < 0.001, η^2^ = 0.579). This finding suggests that cueing distractors with variable predictive validity influence subsequent target-related performance. Planned pairwise comparisons for ES (Fig. S3A) again showed a prominent distractor suppression effect in high predictive validity trials (High minus Null: t25 = 2.501, p = 0.019, two-tailed, Cohen’s d = 0.492) and low predictive validity trials (Low minus Null: t25 = 3.467, p = 0.001, two-tailed, Cohen’s d = 0.680). For the slope of ES (Fig. S3B), a similar distractor suppression effect was observed in the high predictive (High minus Null: t25 = 7.192, p < 0.001, two-tailed, Cohen’s d = 1.411) and low predictive validity trials (Low minus Null: t25 = 6.703, p < 0.001, two-tailed, Cohen’s d = 1.315). However, no significant distractor suppression effect (High minus Low) was observed for ES (t25 = 0.966, p = 0.342, two-tailed, Cohen’s d = 0.190) or slope of ES (t25 = 1.171, p = 0.252, two-tailed, Cohen’s d = 0.230). These results may be due to a ceiling effect of behavioral responses or limitations of current testing paradigms for examining “responses for target” in distractor-related manipulation.

#### Alpha MI of distractor cueing

The alpha MI of high predictive validity (red line) and low predictive validity (blue line) trials during the cue period are shown in Fig. 4B. Due to the cue display without lateralized spatial information, we did not analyze alpha MI in null predictive validity trials. Our results showed that a significant negative alpha MI occurred only in high predictive validity trials during the late period of 878 – 1148 ms, which suggested the alpha power increased in the contralateral hemisphere to distractor cue (permutation test: p < 0.050, two-tailed). Post hoc analysis revealed that alpha MI in high predictive validity trials was significantly lower than that in low predictive validity trials (High minus Low: 867-1113 ms; permutation test: p < 0.050, two-tailed).

#### Distractor-elicited PD

We anticipated that as the predictive validity of the distractor cue increased, the participant’s reactive suppression of the subsequent salient distractor in the search array would decrease, resulting in a smaller distractor-elicited PD. As expected, the results (Fig. 4C) showed a significant main effect of predictive validity on P_D_ (F2, 50 = 3.173, p = 0.049, η^2^ = 0.099). Further paired t-tests confirmed that the P_D_ elicited by expected distractors in high predictive validity trials was greatly reduced in amplitude compared to expected distractors in low predictive validity trials (t25 = −2.126, p = 0.042, Cohen’s d = −0.388) and unexpected distractors in null predictive validity trials (t25 = −2.266, p = 0.031, Cohen’s d = −0.414). The distractor-elicited P_D_ did not differ between low- and null-predictive validity trials (ps > 0.050, BF10 < 0.333).

#### Correlation analysis between alpha modulation and distractor-elicited PD

We used correlation analysis to investigate the relationship between cue-induced alpha lateralization and subsequent distractor-elicited P_D_ in high- and low-predictive validity trials.

When predictive validity of the distractor cue was high, we found a significant correlation between negative alpha MI (averaged 800 – 1100 ms) and P_D_ amplitude (Fig. 5A, left; r = 0.410, p = 0.041), which suggested that subjects with more alpha power contralateral to the cued distractor (negative alpha MI) during the cue-distractor period showed smaller distractor-elicited P_D_ amplitude in subsequent visual searches. However, there was no significant correlation between alpha MI and P_D_ amplitude (Fig. 5C, right; r = 0.025, p = 0.909) when the predictive validity of the cueing distractor was relatively low.

**Fig. 5.**
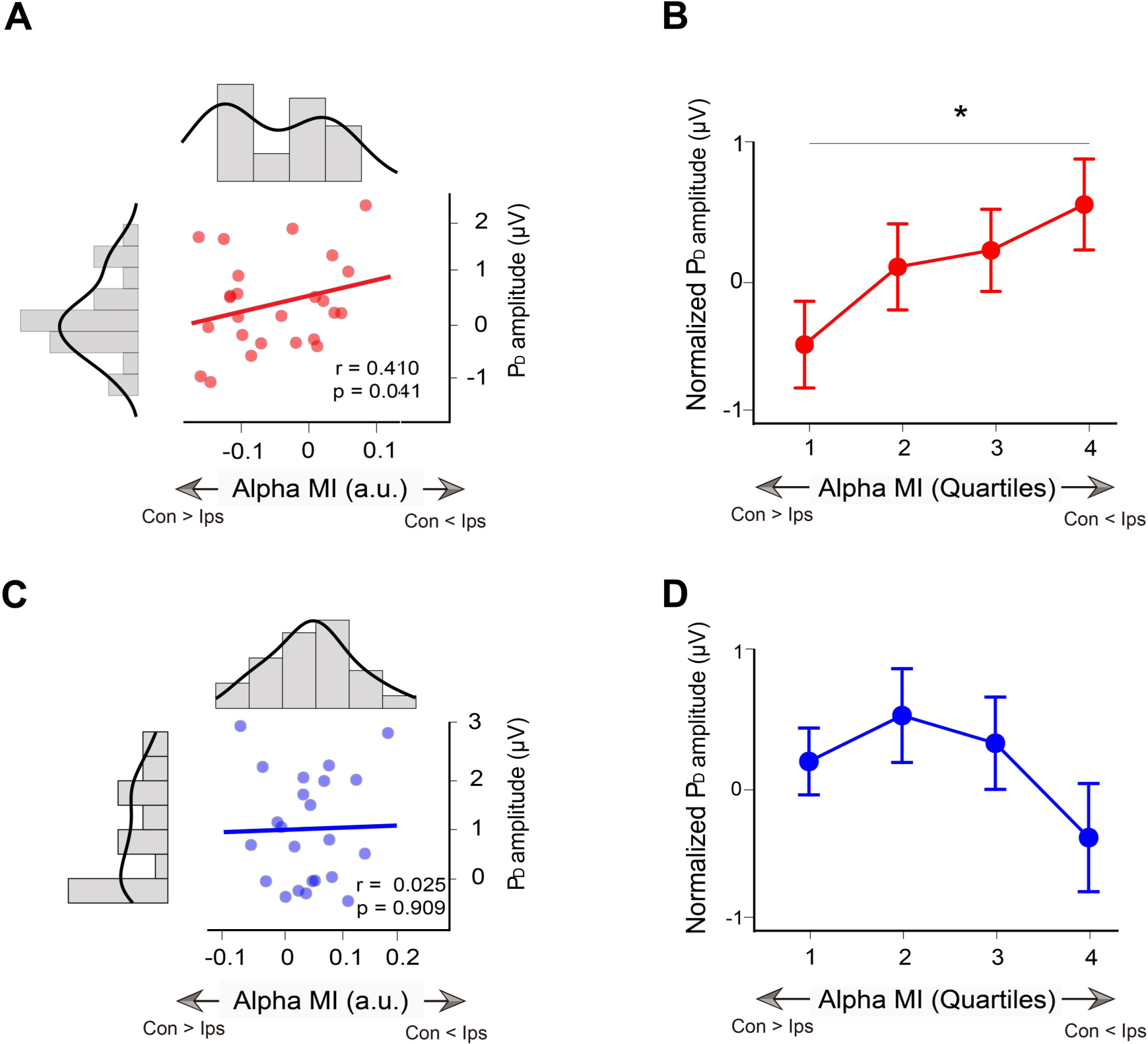
Relationship between alpha MI and P_D_ in Experiment 2. **A** Alpha MI during the cue period as a function of the subsequent distractor-elicited P_D_ amplitudes during a visual search between participants in high-predictive validity trials. The diagrams along with the scatter plot are the frequency distributions of alpha MI and P_D_ amplitude, respectively. **B** Averaged single-trial P_D_ for each quartile at the within-subjects level in high-predictive validity trials. The trials were sorted according to cue-induced alpha MI and binned into quartiles. The P_D_ amplitudes were normalized and then averaged over subjects. *p < 0.05. **C** Scatter plot for low-predictive validity trials. **D** Quartile plot for low-predictive validity trials. Con: contralateral to distractor cue; Ips: ipsilateral to distractor cue.

Furthermore, we calculated the average single-trial P_D_ for each quartile at the within-subjects level. Although the repeated-measures ANOVA of P_D_ did not reach significance (F3, 75 = 1.818, p = 0.152), the results in the high predictive validity trials show that the P_D_ amplitude in the fourth negative quartile was significantly larger than that in the first negative quartile (Fig. 5B, left; t25 = 2.303, p = 0.030, Cohen’s d = 0.461). Accordingly, we suggested that the trials with more alpha power contralateral to the cued distractor (negative alpha MI) also showed smaller distractor-elicited P_D_ amplitude subsequently. Similarly, no significant difference was found among the quartiles (Fig. 5D, right; ps > 0.050) in the low-predictive validity trials. These results showed that there was a close relationship between alpha MI and subsequent biomarkers of distractor suppression at both the between- and within-subjects levels when the predictive validity of distractor cues was high. Behaviorally, we further investigated whether the slope of ES is correlated with the alpha MI or the amplitude of P_D_ in Experiment 2, no correlation was found in different trials (ps > 0.451).

### Experiment 3

To date, the evidence for distractor processes was confined to tasks with a graphic cue (circular radar-like cues) and a modest sample size. The former may cause limited generality, while the latter can increase the false-positive rate and give rise to inflated effect sizes (Yarkoni et al., 2009). Thus, the purpose of Experiment 3 was twofold: 1) further investigate the alpha power modulation of the distractor cue by using the arrow cue to rule out any graph-specific effects and 2) to explore the potential relationship between distractor anticipation and subsequent distractor inhibition based on large sample size (N > 40). The same analysis pipeline as Experiment 2 was applied in Experiment 3.

As shown in Fig. 6A, we again isolated significant negative alpha MI (8-12 Hz) for distractor cues during late cue-distractor intervals (permutation test: p < 0.050, two-tailed). Grand averaged ERPs locked to distractor onset were generated to calculate the P_D_ component. The P_D_ was significantly different than zero from 234 to 330 ms (Fig. 6C, permutation test: p < 0.050, two-tailed). The scatter plot showed a significant correlation between alpha MI (averaged 750–950 ms) and P_D_ amplitude (Fig. 6D, r = 0.332, p = 0.028). We suggest that subjects with a more alpha power contralateral to the cued distractor have a lower P_D_ amplitude in subsequent visual search.

**Fig. 6.**
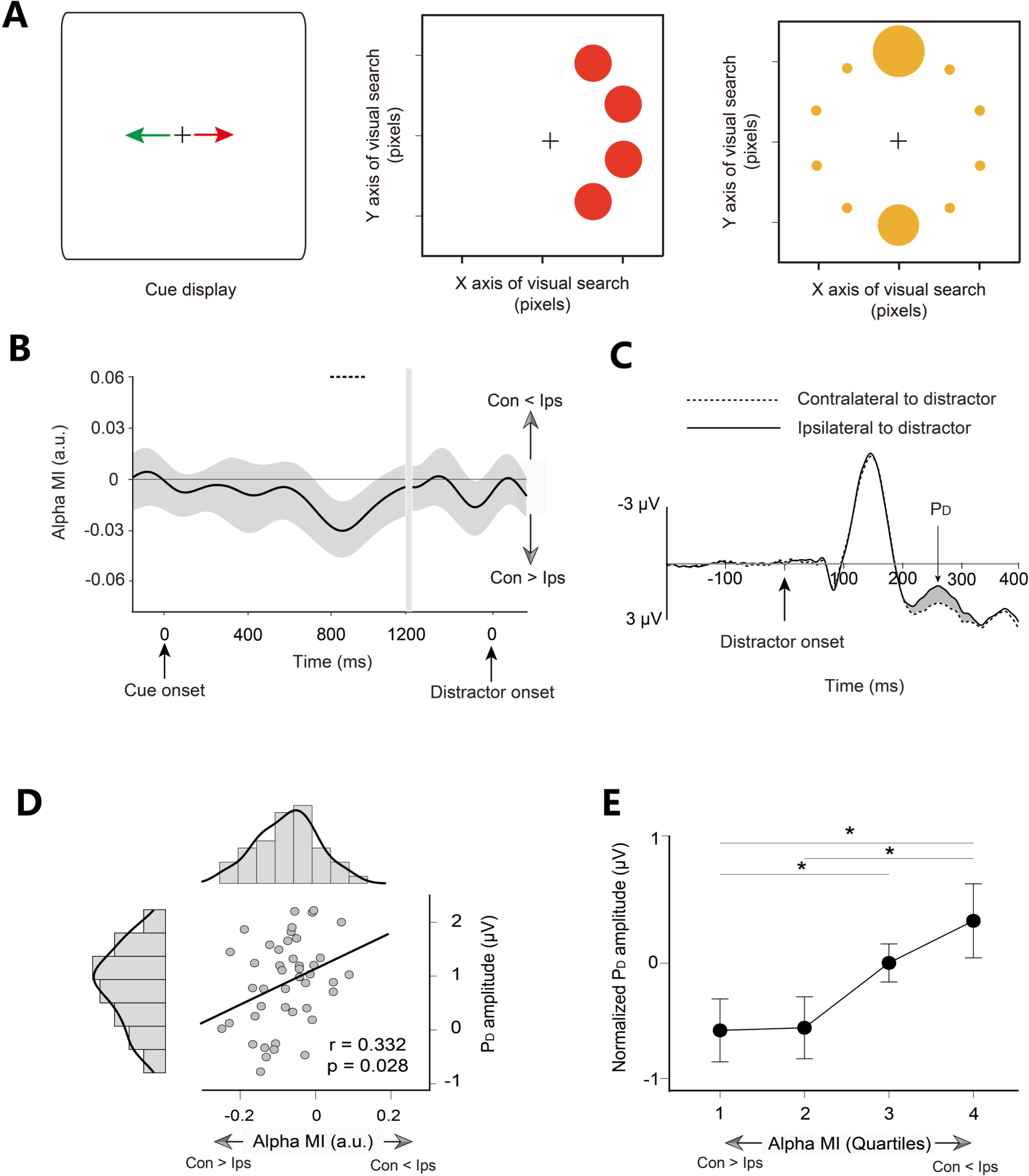
Task paradigm and EEG results for Experiment 3. **A** The arrow was fully predictive of the side on which the distractor circle of the corresponding color would subsequently appear. **B** Time course of the alpha MI. **C** Grand averaged ERPs at contralateral and ipsilateral electrode sites relative to the distractor. **D** The scatter plot between cue-induced alpha MI (averaged over the time-frequency windows highlighted by black outlines) and distractor-elicited P_D_ amplitudes between participants showed a significant correlation. The diagrams along with the scatter plot are the frequency distributions of alpha MI and P_D_ amplitude, respectively. **E** Averaged single-trial P_D_ for each quartile at the within-subjects level. Trials were sorted according to cue-induced alpha MI and binned into quartiles. P_D_ amplitudes were normalized and then averaged over subjects. *p < 0.05. Con: contralateral to distractor cue; Ips: ipsilateral to distractor cue.

Here, we also calculated the average single-trial P_D_ for each quartile at the within-subjects level by the same method as applied in Experiment 2. We found that the alpha MI induced by spatial cues strongly correlated with the subsequent P_D_ component: the normalized P_D_ amplitude decreased with an increase in alpha power contralateral to the cued distractor (Fig. 6E, repeated-measures ANOVA, F3, 123 = 3.078, p = 0.030). Simple first contrast shows that the P_D_ amplitude in the fourth negative quartile was significantly larger than that in the first (t41 = 2.059, p = 0.046, Cohen’s d = 0.330) and second negative quartiles (t41 = 2.171, p = 0.036, Cohen’s d = 0.348). The P_D_ amplitude in the third negative quartile was significantly higher than that in the first negative quartile (t41 = 2.184, p = 0.035,

Cohen’s d = 0.350), suggesting that the trials with more alpha power contralateral to the cued distractor during the cue-distractor period have a less distractor-elicited P_D_ amplitude in the following visual search. In sum, a close relationship between alpha lateralization and subsequent biomarkers of distractor suppression was further confirmed between and within subjects in Experiment 3.

## Discussion(1500)

The current study gains more insight into the neural mechanisms underlying proactive suppression guided by spatial cues. Across three experiments, we presented a series of evidence on the existence of alpha activity related to proactive suppression and how it shapes subsequent distractor processing. In Experiment 1, a cueing distractor could sharpen the spatial behavioral measurement (slope of ES), induce distractor suppression (negative alpha CTF slope), and reduce distractor interference (PD amplitude). The analysis further showed that negative alpha CTF was related to reduced distractor-elicited PD. Results from Experiment 2 demonstrated that increased alpha power contralateral to the distractor (negative alpha MI) and the reduced P_D_ may be the result of distractor suppression derived from spatial effectiveness. Crucially, when spatial cues with high predictive validity were employed, the P_D_ amplitude was observed as a function of cue-elicited alpha MI. That is, a more increased alpha power contralateral to the cued distractor (negative alpha MI) has less distractor interference, as reflected in the decreased PD. Additionally, a symbolic cue with high predictive validity was further employed in Experiment 3. The significant correlation across individuals and quartile analysis within individuals was further confirmed. Together, these results have shown how spatial distractor foreknowledge proactively reduces distractor interference.

Behaviorally, Fig. 1C shows no difference in mean measurements between valid and invalid sessions, which was consistent with previous studies (Wang et al., 2018). Interestingly, we found a dichotomous phenomenon in Experiment 1: cueing distractors precisely had a trend towards harming the performance on the target that appeared in close spatial proximity of the distractor, but boosting the performance on the target appeared in far away spatial proximity of the distractor. To measure this trend, we showed spatial changes as a function of the distances of the target to the distractor and found a significant distractor cueing effect (valid - invalid) on spatial changes. In Experiment 2, we further found that such spatial changes were related to the spatial effectiveness of cueing distractors. Our results go beyond previous research by demonstrating that the performance of the target was influenced not only by statistical learning (Wang et al., 2018) but also by the spatial intention of the distractor.

From distinct research traditions, alpha CTF (Fig. 2C) and alpha MI (Fig. 2E) provided convergent evidence for the dynamic characteristic of proactive suppression for Experiment 1. In the beginning, regardless of its task relevance, spatial cues with spatial information inputs may result in increased distractor selectivity (positive alpha CTF slope) or relatively decreased alpha power contralateral to cue distractor (positive alpha MI) at the early stages. This result was consistent with Foster’s (2017a) findings and suggests that enhanced tuning towards cued directions might be first represented in our spatial attention. Then, we observed that our brain engages the progressive attenuation of both the amplitude of alpha MI and CTF slope. We suggest that this result stems from a white-bear metaphor (Tsal & Makovski, 2006), in which participants have to make an effort to minimize interference from the distractor per se when given distractor information. Interestingly, during the late preparatory stages, our results clearly show the distractor suppression (negative alpha CTF slope or negative alpha MI) in valid sessions. The results from Experiment 2 and Experiment 3 further confirmed the existence of such alpha activity through the negative alpha MI.

Based on the negative CTF slope result in Fig. 2, alpha MI during the late period of cue-distractor intervals (see Fig. 2, 4, and 6) could also be considered the result of the alpha CTF in the case of lateral cues. As an example, in Experiment 1 (Fig. S5), when the cue pointed left (e.g., θ = 288 °), the channel response (alpha power) decreased over the left hemisphere and increased over the right hemisphere; when the cue pointed right (θ = 108°), the channel response (alpha power) increased over the left hemisphere and decreased over the right hemisphere. We calculated such an asymmetric channel response (alpha power) by collapsing across attend-left and attend-right conditions and collapsing across hemispheres (see Methods for more details). The lateralized channel response and observed lateralized alpha power have similar dynamics (compare Fig. 2C and 2E) and spatial patterns (Fig. S5C). Thus, we inferred that the negative CTF slope might provide a general computational model for the negative alpha MI observed in our study. However, the relationship between CTF and alpha MI needs further study.

A recent study (van Moorselaar et al., 2020) outlined three potential computational models for accounting for distractor suppression within the CTF framework. This suggests that distractor-related negative tuning may arise as a consequence of enhanced tuning towards the opposite distractor direction, shifting sensory tuning away from the distractor direction, or a combination of both. Through comparison with invalid-cue sessions, our results suggest that distractor suppression might result in both tuning towards the opposite distractor direction and away from the cued distractor direction (Fig. 2D), which fits well with the interpretations of the above the third models. Based on this model, alpha power increases at electrodes far away from the cued distractor and decreases at electrodes close to the cued distractor, so that the to-be-captured resources would be relatively diminished from distractors to support target-related activities. We suggest that during cue-distractor intervals, a template-to-distractor (or spatial priority map) might be architected by the gating role of alpha activity

By comparing P_D_ amplitude in Experiment 1, we found that cueing distractors are likely to reduce the amplitude of PD. This result was consistent with van Moorselaar’s study (2019), in which distractor expectations reduced distractor-specific processing, as reflected in the disappearance of PD. Our results in Experiment 2 further expand this idea and suggest that reduced P_D_ was not only related to whether the cue was effective or not but also related to whether the predictive validity of the distractor was effective (Fig. 4C). Crucially, the correlation across subjects and quartile analysis further showed that reduced P_D_ amplitude was a function of alpha MI. That is, the more alpha power contralateral to the cued distractor is, the lower the P_D_ amplitude is. Given that a reduced P_D_ is correlated with minimized distractor interference (Liesefeld et al., 2017), we argue that the brain can engage in proactive filtering mechanisms that operate attention resources that are less likely to be deployed to a cued distractor, resulting in less interference by the subsequent distractor.

Note that such transient modulation of alpha power and its link with P_D_ amplitude does not occur throughout the anticipation period — until the presentation of the search display. One possible explanation is that the participants might strategically have no incentive to persist in suppressing the direction of the task-irrelevant distractor in advance, especially at the cost of task-relevant targets likely occurring in the nearby cued direction. We also suggest another possible explanation that participants seem able to proactively suppress distractors at a cued location by nonconsecutive alpha modulation. Given that visual and memory systems are reciprocally connected (Awh et al., 2001; Gazzaley et al., 2012; Forster et al., 2018), alpha power lateralization also reflects spatial inhibition processes stored in working memory (Rösner et al., 2020), we suspected that a template-to-distractor might not be persistent until the onset of the search display. Alternatively, it was temporarily stored in a visuospatial sketchpad. Once the onset of the visual search was detected, the template-to-distractor can be used to suppress the distractors without feedforward communication of distractor information involving reactive suppression (Geng et al., 2014). Our ERP results supported the above hypothesis by showing that no significant P_D_ followed after a significant negative alpha MI in Experiments 1 and 2. This seems to mean that distractors can be directly suppressed at the low neural level (posterior cortex) in the early stage (∼200 ms), resulting in the null of distractor-elicited P_D_ (approximately 200∼ ms). Importantly, the significant relationship between transient alpha modulation and P_D_ amplitude might provide meaningful evidence for the above hypothesis. However, our results showed that significant P_D_ followed after a significant negative alpha MI in Experiment 3, the absence of a P_D_ effect and its link with alpha activity should be interpreted with caution, and further studies are necessary to gain a better understanding of the template-to-distractor that plays a key role in distractor suppression.

In summary, our results show a series of unambiguous evidence for the underlying neural mechanism of proactive suppression, in which alpha power plays an important role in reducing distractor interference when it appears. In our study, proactive suppression relies on dynamic intentions guided by spatial cues in different circumstances and presents alpha activity in different guises (CTFs or alpha MI). Importantly, a strong link between cue-elicited alpha power and distractor-elicited P_D_ suggests that alpha power activity may reduce interference following distractor onset. These findings contribute to the growing body of work showing that distractor suppression is flexible and involved in more than one general top-down mechanism (Noonan et al., 2018; Geng et al., 2019; van Moorselaar et al., 2020). (1488)

## Materials and methods

### EEG recording and preprocessing

In all experiments, continuous EEG was recorded using a SynAmps EEG amplifier and Scan 4.5 package (NeuroScan, Inc.). In Experiment 1, EEG data were recorded from 15 international 10-20 sites, F3, Fz, F4, T3, C3, Cz, C4, T4, P3, Pz, P4, T5, T6, O1, and O2, along with five nonstandard sites: OL midway between T5 and O1, OR midway between T6 and O2, PO3 midway between P3 and OL, PO4 midway between P4 and OR, and POz midway between PO3 and PO4. In Experiments 2 and 3, EEG data were recorded using a 32-electrode elastic cap (Greentek Pty. Ltd) with silver chloride electrodes placed according to the 10-20 system. To detect eye movements and blinks, horizontal electrooculograms (HEOG) and vertical electrooculograms (VEOG) were recorded via external electrodes placed at the canthi of both eyes, above and below the right eye, respectively. All electrodes, except those for monitoring eye movements, were referenced to the left mastoid during data collection and then were off-line re-referenced to the algebraic average of the left and right mastoids. The EEG was amplified with DC-200 Hz, digitized on-line at a sampling rate of 1000 Hz (sampling interval 1 ms), and then off-line filtered with a digital bandpass of 0.1–40 Hz (6 dB/octave roll-off, FIR filter). We kept electrode impedance values below 5 kΩ.

EEG data were preprocessed using the EEGLAB software package in the MATLAB environment (Delorme and Makeig, 2004). Independent component analysis (ICA, EEGLAB runica function) was performed for continuous data. Component removal was restricted to blink artifacts (less than two on average).

Trials in which the EEG exceeded ±100 μV in any channel and the horizontal EOG exceeded ±50 μV from −200 to 400 ms in the cue- or distractor-locked epochs were automatically excluded in all experiments. Overall, artifacts led to an average rejection rate of 15.4% of trials (range 7.1–23.7%) in Experiment 1, 18.0% (range 11.2–31.7%) of trials in Experiment 2, and 17.9% of trials (range 8.2–29.1%) in Experiment 3. A total of 857 (SD: 49) for each session in Experiment 1, 264 (SD: 28) for each trial in Experiment 2, and 273 (SD: 28) in Experiment 3 were used for further analyses.

### Inverted encoding model analysis

For the inverted encoding model (IEM) analysis, we followed a similar approach to the previous work (Foster et al., 2017b). We used an IEM to reconstruct location-selective CTFs from the topographic distribution of EEG activity across electrodes to examine the spatially specific alpha-band activity time course. Briefly, this model assumes that the power at each electrode (one per sample angle) reflects the weighted sum of ten spatially selective channels (Brouwer & Heeger, 2009; Sprague & Serences, 2013). We modeled the responses of each electrode using a basis function of ten half-sinusoids raised to the ninth power for each spatial channel: R = sin(0.5θ)^9^, such that θ is the angular location (0°, 36°, 72°, 108°, 144°, 180°, 216°, 252°, 288°, 324°) and R is the spatial channel response.

EEG data were segmented into 2000 ms epochs ranging from 500 ms before to 1500 ms after cue onset for the cue-locked analysis. Data were also segmented and aligned according to target onset from −800 to 800 ms for the distractor-locked analysis. Then, EEG segments were bandpass filtered for the alpha band (8-12 Hz) using a function (eegfilt) from the EEGLAB toolbox (Delorme & Makeig, 2004). The filtered data were transformed to instantaneous power using a function (Hilbert) from MATLAB (The Mathworks, Natick, MA). The IEM was run on each time point in the alpha band power.

We sorted the artifact-free trials into training sets (B1) and test sets (B2) for each subject (for details, see Foster et al., 2017b). Let B1 and B2 be the power at each electrode for each trial in the training set and test set, respectively. Data from the training set (B1) were used to estimate channel-to-electrode weights on the hypothetical spatial channels separately for each electrode. The basis functions determined the channel response function (C1) for each spatial channel.

The training data (B1) in electrode space were then mapped onto the matrix of channel outputs (C1) in channel space by the channel-to-electrode weight matrix (W), which was estimated with a general linear model of the form:

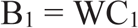

The estimated channel-to-electrode weight matrix can be derived via least-squares estimation as follows:

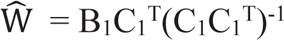

In the test stage, channel responses (C_2_) were estimated based on the observed test data (B2) with the weight matrix W:

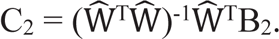

Finally, the ten estimated response functions (C2) were aligned to a common center. The center channel was the channel tuned for the location of the specific stimulus (i.e., θ°) and then averaged to obtain the CTF. The CTF slope was used as a metric to compare attention deployment towards the distractor.

### Alpha modulation analysis

The segmented EEG data were decomposed using Morlet wavelet-based analysis from 8 to 12 Hz in 1 Hz steps implemented in the related package Brainstorm (Tadel et al., 2011) in the MATLAB environment. We subtracted the trial-average activity in the time domain from the EEG activity of every single trial to avoid the time-frequency power being disturbed by the ERP in oscillatory signals.

To estimate the effects of cue-elicited attention modulation, we calculated the alpha modulation index (MI) from cue-locked data for three pairs of parietal and occipital electrodes (left ROI: P3, P7, O1; right ROI: P4, P8, O2). The MI was computed using the following formula:

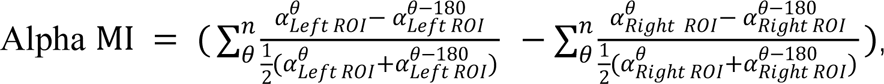

where θ indicates the angle of cue pointing (θ = 288° or 252° in Experiments 1; θ = 270° in Experiments 2 and 3); α indicates alpha band power within the left ROI or right ROI; and n ^is the number of θ in the modulation analysis (n =2 in Experiment 1; n = 1 in Experiments 2^ and 3).

Note that the above method allowed us to avoid possible bias in the analysis due to the hemisphere asymmetry (Zhao et al., 2022). The amplitude of MI denotes the deviation of spatial alpha power in the hemisphere contralateral to the cued distractor with respect to the hemisphere ipsilateral to the distractor. Further, the polarity of alpha MI denotes the direction of spatial alpha modulation: positive values indicate alpha power relatively decreases contralateral to distractor; negative values indicate alpha power relatively increases contralateral to the distractor. We also plotted alpha power in contralateral and ipsilateral to the cued distractor respectively across three experiments (see Fig.S6 for the results).

### Decoding analysis

We adopted the same procedure as reported in a previous study (van Moorselaar et al., 2020), except with 22 EEG channels as features and the spatial location of distractors or targets as classes. In brief, we used multivariate pattern analysis (MVPA) in combination with linear discriminant analysis to assess whether the spatial distribution of EEG data could be used to decode the distractor or target location in Experiment 1. The performance of decoding based on EEG data is the 10-fold cross-validation AUC (area under the ROC curve) of the corresponding model.

### ERP analysis

The EEGLAB toolbox (Delorme & Makeig, 2004) and ERPLAB toolbox (Lopez-Calderon & Luck, 2014) were used to process and analyze ERP. The combination of a lateral distractor and a midline target (see Fig. 3) enables the isolation of EEG activity in response to the distractor (Gaspar et al., 2014). Thus, we analyzed the ERP elicited by the subsequent visual search display with a lateral distractor and midline target to isolate distractor-specific P_D_ components. ERP was computed by subtracting the waveforms measured from electrodes (P7 or P8) on the ipsilateral hemisphere to the distractor from symmetrical electrodes on the contralateral hemisphere. Then, ERP was corrected using a −200 to 0 ms window preceding stimulus onset. Finally, the amplitude of P_D_ was achieved in the ERPLAB measurement tool as the mean value of a 20-ms window centered at the most positive peak in the averaged difference waveform between 220 ms and 320 ms.

### Correlation and quartile analysis

We performed a similar time-frequency correlation method reported in Zhao et al. (2019). We extracted alpha MI values based on a 60 ms sliding time window (steps of 5 ms) across a time range of −200 to 1200 ms for each subject and then correlated them with distractor-evoked PD. Each pixel of the time-frequency correlation map consisted of Pearson’s r value between alpha MI at each time interval and each frequency and subsequent P_D_ amplitude. Then, the significant spectrogram related to PD amplitude (p < 0.050) was corrected for false-discovery rates (FDR) within a prior defined frequency range of 8-12 Hz across the entire time. The left significant spectrogram (P_corrected_, < 0.050) was defined as the TFC ROI.

We also adopted a similar quartile analysis within subjects as reported in Dijk et al. (2008). The average single-trial P_D_ was estimated at the within-subjects level to confirm the relationship between the alpha MI and subsequent P_D_ amplitude. The trials were sorted according to alpha MI and split into quartiles for the attend-left session and attend-right session. The separate P_D_ waveforms for each session were calculated for each quartile and normalized to the individual mean value over all quartiles. The final P_D_ for each quartile was computed by averaging the P_D_ from the right- and left-attend sessions.

### Participants

One hundred and ten paid volunteers participated in the three experiments (Experiment 1: 32, Experiment 2: 28, Experiment 3: 50), twelve of whom were excluded from statistical analysis due to excessive EEG artifacts (2 participants in Experiment 1, 2 participants in Experiment 2, 8 participants in Experiment 3). Data from the remaining 30 participants in Experiment 1 (18 male, 22.6 years mean age), 26 participants in Experiment 2 (17 male, 22.7 years mean age), and 42 participants in Experiment 3 (30 male, 23.2 years mean age) were used. All participants had a normal or corrected-to-normal vision and were right-handed. They were neurologically unimpaired and gave informed written consent before the experiment. All experiments were conducted in accordance with the Beijing Normal University Institutional Review Board.

### Task, stimuli, and procedure

Previous studies (van Moorselaar et al., 2020; Wang et al., 2018) have arranged target and distractor locations by dividing 2D space into four or six parts. Similar to black and white, spatial distractor cues might indirectly provide potential spatial information about a target, e.g., when the distractor was occurring on the left, the target was presented on the right more often, and vice versa. Considering that participants pick up such statistical regularities and use them to guide their target selection (Geng et al., 2005; Wang et al., 2018), increased alpha power contralateral to the distractor might be mixed by potential target-related activity (decreased alpha power contralateral to more often the target). Thus, ensuring that participants do not have target-related activity is essential to study distractor suppression, which is also in compliance with the relevant principles (see rule 2 in Wöstmann et al., 2022). In this sense, we minimized target-dependent activity by increasing the number of possible directions (N=10) and decreasing the probability of the target occurring on the lateral side (Experiments 2, 3).

In this study, three experiments were conducted to investigate the influences of the spatial cues of the distractor on the subsequent visual search. In each experiment, a 200 ms cue informed the participants of the location (Experiment 1) or scope (Experiment 2, 3) in which the upcoming distractor would occur in the search display. The cue-distractor interval was 1200–1600 ms. Each search display consisted of 10 unfilled circles presented for 200 ms (13.5 cd/m^2^ mean optical luminance, and 3.4°×3.4°, 0.3° thick outline) from the imaginary ring with a 9.2° radius. A yellow target circle and a red distractor circle were simultaneously presented among the eight green circles. A schematic of the trial design is illustrated in Fig. 1A.

Salience was defined in terms of the local contrast between green circles and each color circle (see Fig. S5): the distance in chromaticity space between the red distractor (RGB: 255, 100, 100) and green circles (RGB: 0, 180, 0) was greater than the distance between the yellow target (RGB: 160, 160, 0) circle and green circles. A red distractor with more salience captures attention more easily than a yellow target, causing more incentive to ignore distracting sensory information. Participants were instructed to utilize the cue to ignore a more salient distractor (red circle) and determine whether the line segment inside the target (yellow circle) was vertical or horizontal by pressing one of two buttons with their right hand as quickly as possible.

#### Experiment 1

In Experiment 1, a red circular sector with an angle of 36° was embedded in a full green circle at the center of the display (see Fig. 1A), which randomly and equally pointed to one of ten possible directions (0 °, 36 °, 72 °, 108 °, 144 °, 180 °, 216 °, 252 °, 288 °, or 324 °) with reference to the upper y-axis (0 °). As shown in Fig. 1B, this graphic cue was typically informative for the valid-cue session (100% probability on a cued direction) or uninformative for the invalid-cue session (10% probability on a cued location) of the location at which the subsequent red distractor circle emerged. In both valid- and invalid-cue sessions, the location of the subsequent target was independent of which distractor location and randomized with equal probability (10% probability on each location), so that subjects could not infer anything about the yellow target circle from the cue. The sequence of the two sessions was counterbalanced between subjects. Each session consisted of ten 100-trial blocks and lasted approximately 60 minutes. Participants came to the lab twice, separated by one week.

#### Experiment 2

There were three kinds of graphic cues in Experiment 2. The circular sector was equally likely to point left (90 °) or right (270 °), and the variable area of the circular sector was related to the predictive validity of distractor occurrence. As shown in spatial probability in Fig. 4A, (1) in the high predictive validity trials, the red sector with a polar angle from 216° to 324° (or from 36° to 144°) was fully predictive with 100% validity for the left (or right) side where the red circle distractor would appear, that is, the distractor would appear randomly on one of the cued lateral locations with 25% probability; (2) in the low predictive validity trials, a red semicircle predicted that the red circle distractor would appear randomly on one of the cued locations (with 16.7% probability on one the lateral location or one midline location); (3) in the null predictive validity trials, none of the red sectors embedded in the green circle was uninformative of the upcoming distractor (10% probability on each location). To isolate the brain activity related to distractor anticipation, we pseudorandomized the location of a yellow target circle by specifying a uniform spatial probability of 4.25% on each lateral location and 33% on each midline location (see Fig. 4B; right panel). The experiment contained 10 blocks (i.e., 100 trials per block) per participant. The three types of trials were randomized within each block. Experiment 2 lasted approximately 60 min.

#### Experiment 3

In Experiment 3, we used the constant arrow instead of the variable circular sector as a symbolic spatial cue (see Fig. 6A, left panel). The red arrow was fully predictive of the side (with 100% validity) on which the following red distractor circle would subsequently appear, that is, the distractor would appear randomly on one of the cued lateral locations with 25% probability (Fig. 6A, middle panel). The opposite green arrow had no predictive value for the yellow target circle and red distractor circle. Target had the same spatial probability as that of Experiment 2 (Fig. 6A, right panel). In fact, the cue in Experiment 3 was the same as the high predictive validity trials in Experiment 2 except for the symbolic form of a spatial cue. We called it the arrow high predictive validity cue.

## Notes

### Competing Interest Statement

The authors have declared no competing interest.

