## Supplemental Information for "Neural mechanisms underlying distractor suppression guided by spatial cues"

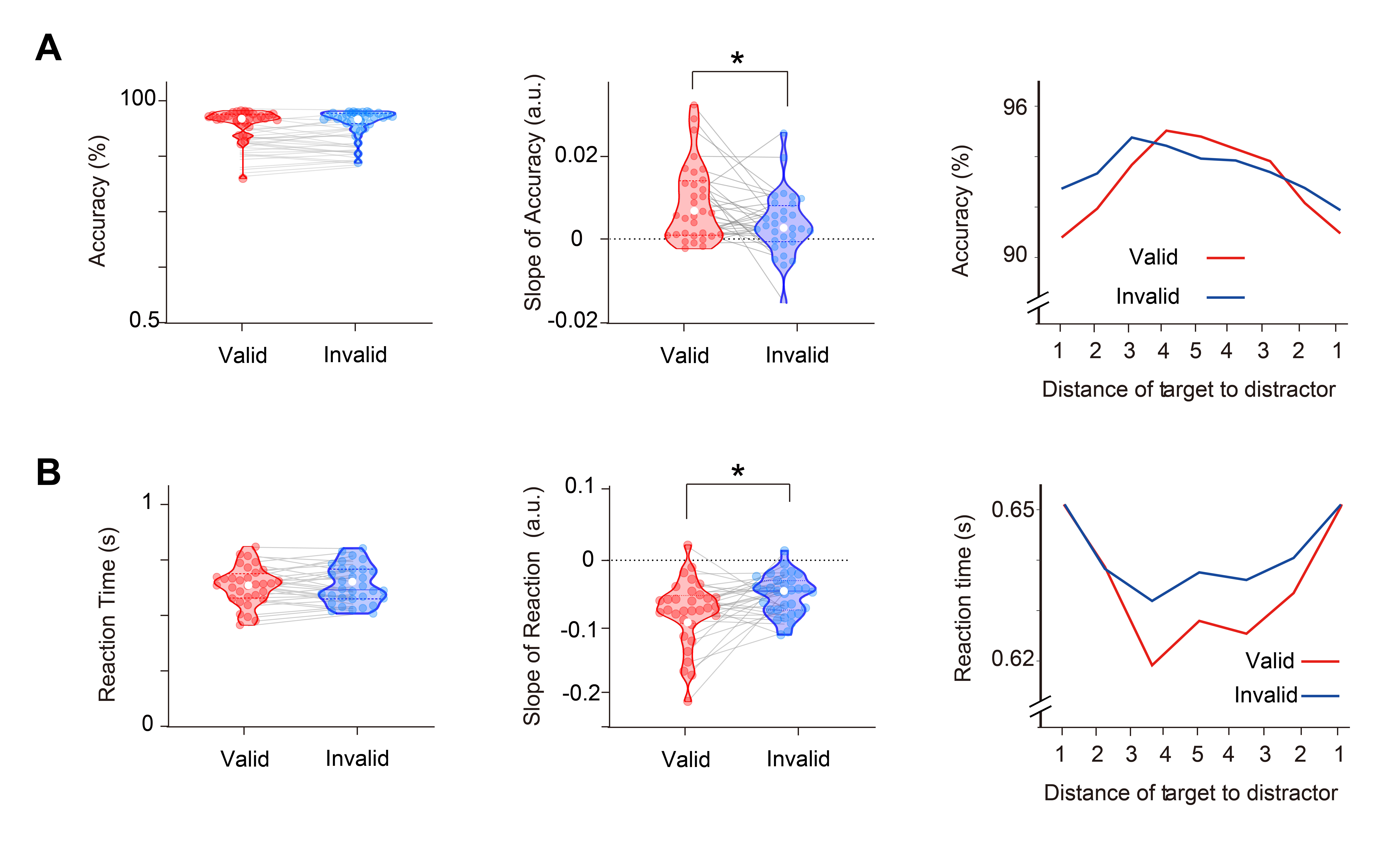

**Fig. S1.** Reaction time and accuracy for Experiment 1. **A** The mean (left) and slope (middle) of accuracy across DTD (right) in the valid (red) and invalid (blue) cue sessions. **B** The mean (left) and slope (middle) of reaction time across DTD (right) in the valid (red) and invalid (blue) cue sessions. Violin plots depict the distributions of measurements in each session, with the dots representing each subject. The solid and dotted lines indicate medians and quartiles, respectively. *p < 0.05.

### *Decoding for distractor and target*

As shown in Fig. S2, AUC (area under ROC curve) scores of distractor decoding were significantly different from the chance level in both valid-cue (142–320 ms, permutation test: p < 0.001) and invalid-cue sessions (154–322 ms, permutation test: p < 0.001). AUC scores for target location also showed a significant difference from chance level in the two sessions (valid: 236–400 ms, permutation test: p < 0.001; invalid: 232–400 ms, permutation test: p < 0.001). The decoding performance for the target was significantly better in valid-cue sessions from 268 to 400 ms (Fig. S2, right panel, permutation test: p < 0.01).

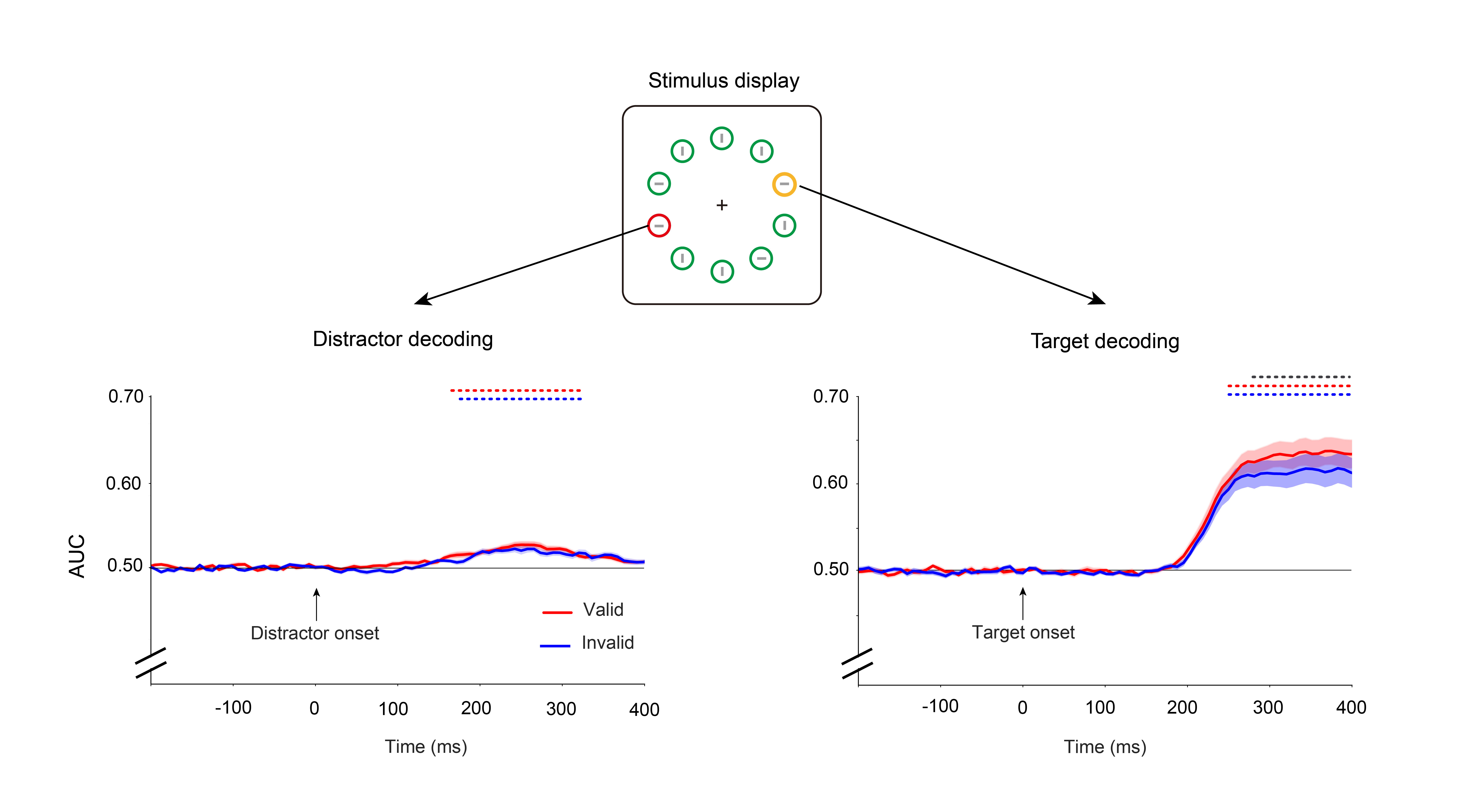

**Fig. S2.** Multivariate decoding during the stimulus period from Experiment 1. AUC scores of distractor (left panel) and target (right panel) spatial location decoding across time in valid (red) and invalid (blue) cue sessions. Shades of light color along with the dark color lines represent error bars (±1 SEM). The red and blue dashed lines at the top indicate clusters where sessions differed significantly from chance after cluster correction (p<0.05), and significant differences between sessions are marked by the black dashed line (p<0.05).

When the spatial cues were predictive, our results showed a better target decoding performance relative to invalid cues (Fig. S3A). These findings may provide further support for the idea that cueing distractors can improve attention allocation to the target (Arita et al., 2012). However, poststimulus distractor decoding performance did not differ. One possible reason may be the low signal-to-noise ratio of distractor-related activity. Previous studies (Wöstmann et al., 2019) have shown that the strength of alpha lateralization for target selection was larger than that for distractor suppression. In practice, effects related to target processing are often considerably larger than effects related to distractor suppression (see rule 9 in Wöstmann et al., 2022). The target decoding performance far outweighs the max value of the distractor (see Fig. S3) based on the same EEG datasets. Similar to winner-takes-all, this suggests that the location and feature of the target will take almost all attentional resources as long as the target appears, which results in an overwhelming target-related activity and easily drowned distractor-related activity. In this sense, the combination of a lateral distractor and a midline target, as shown in Fig. S3B, enables the isolation of lateralized activity in response to the distractor. This allowed us to examine distractor-related processes and avoid mixed target-related activity.

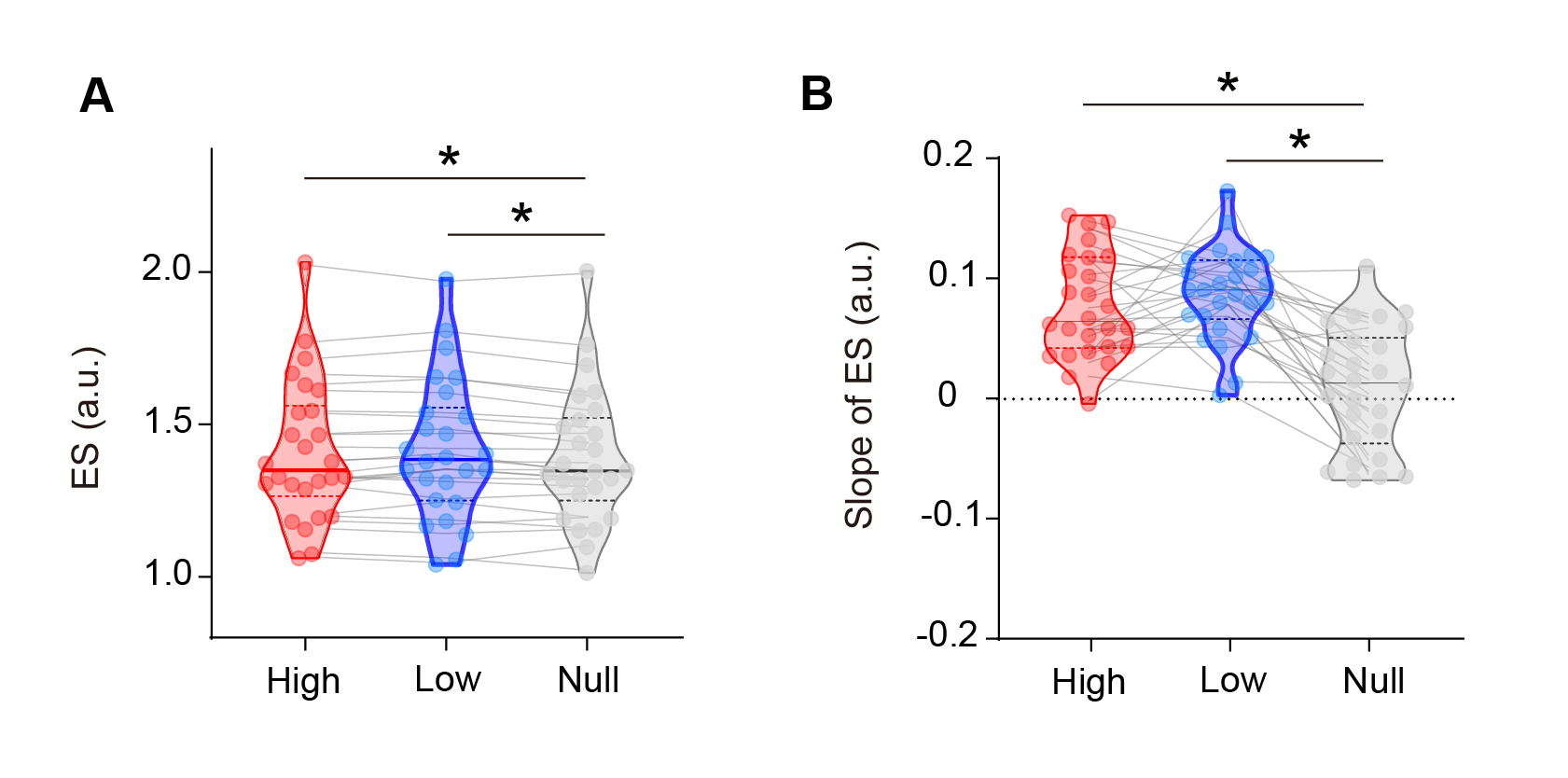

**Fig. S3.** Behavioral results for Experiment 2. **A** Average ES related to the target for high (red), low (blue), and null (black) trials. **B** The slope of ES. The abbreviated letters indicate the three cue display configurations as follows: High, high predictive validity cue; Low, low predictive validity cue; Null, null predictive validity cue. The three cues were randomized within each block. *indicates P < 0.05.

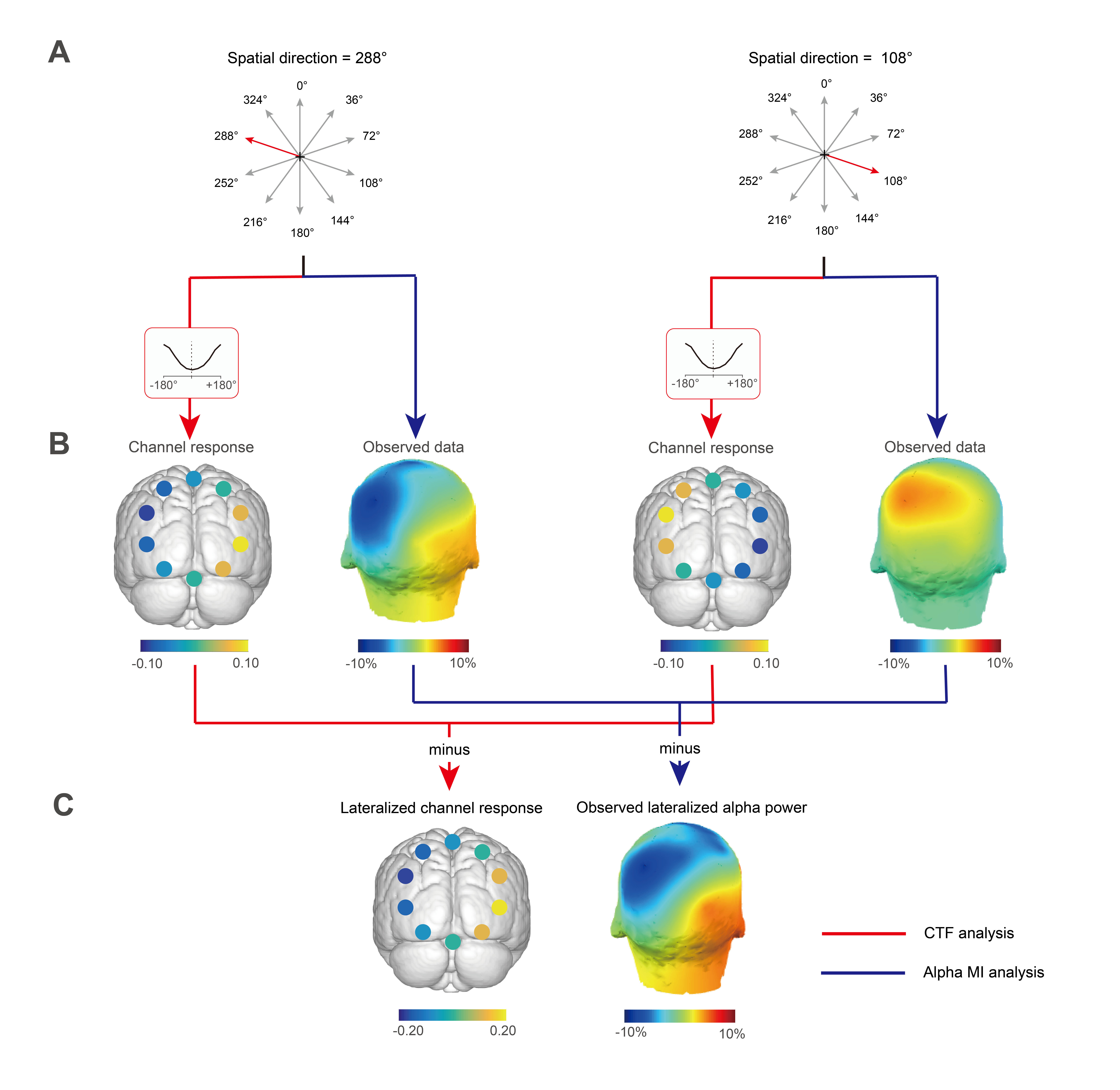

**Fig. S4.** The analysis pipeline for CTF analysis (red line) and alpha MI (blue line). **A** The example shown here is for a pair of spatial directions pointed at 288° and 108°. **B** To show spatial gradient effects related to distractor cues, the spatial change in alpha power was rendered by mapping the channel response curve to ten ideal spatial channels. The scalp topographies show the distribution of alpha power over the posterior cortex. **C** The resulting lateralized channel response and observed lateralized alpha power reflect a similar spatial pattern.

### Fig. S5

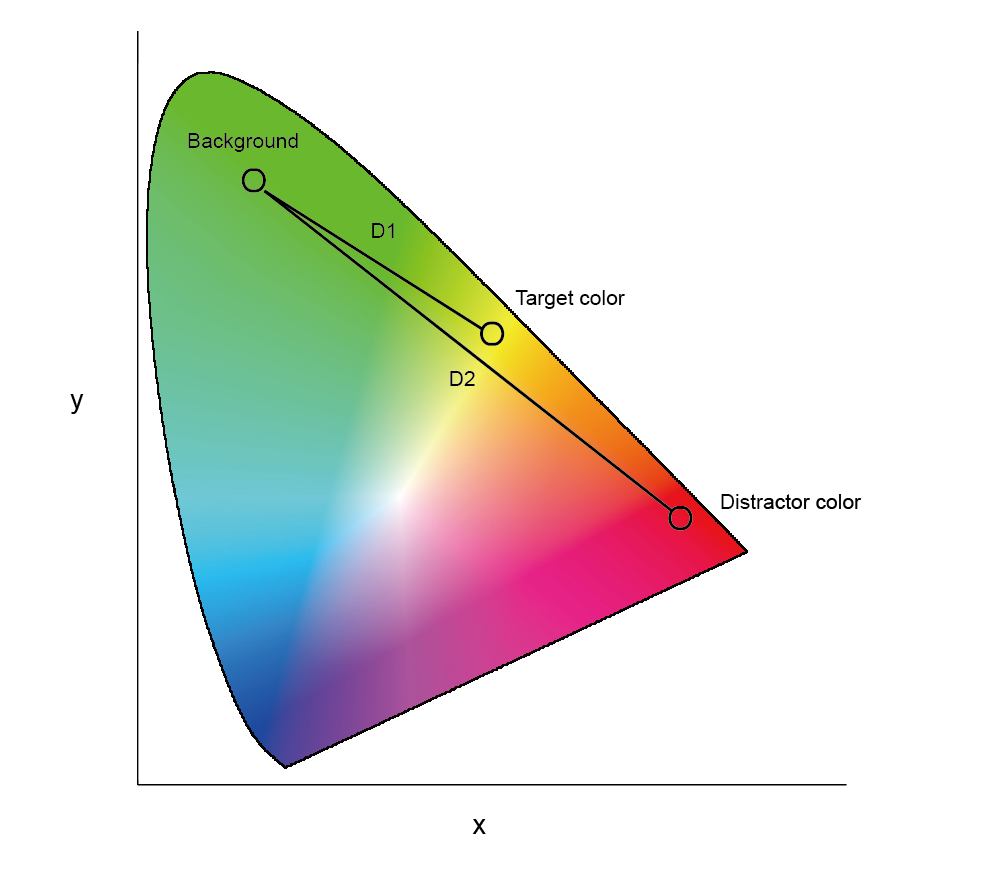

**Fig. S5.** Colour space in the present study. D1: the chromaticity space color distance between the yellow target circle and green circles. D2: the chromaticity space colour distance between the red distractor and green circles.

### Fig. S6

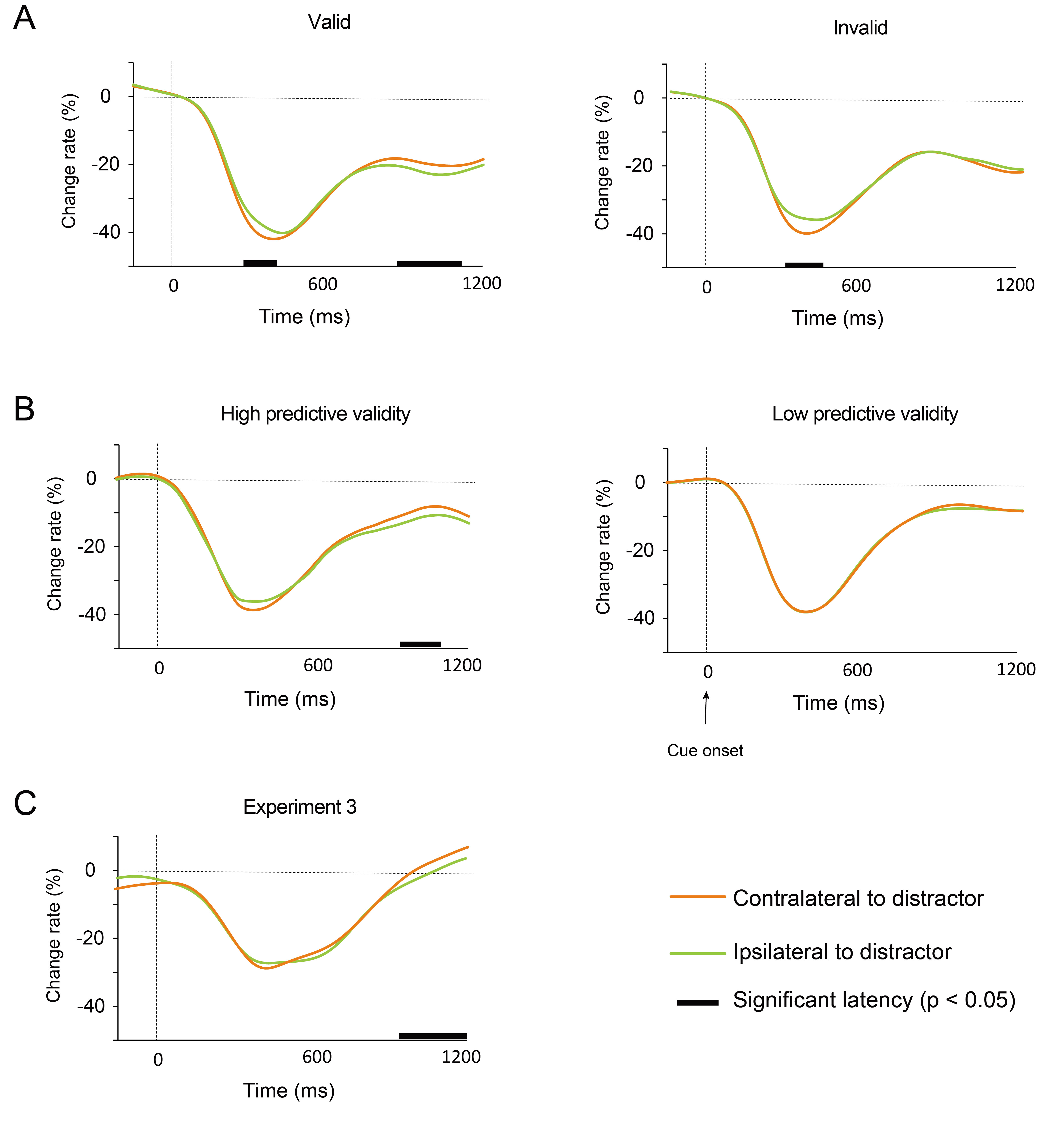

**Fig. S6.** Time course of the change rate of alpha power in contralateral (orange) and ipsilateral (green) hemisphere to distractor cue for Experiment 1 (**A**), Experiment 2 (**B**), and Experiment 3 (**C**). The black lines indicated the change rate of alpha power with a significant difference between the contralateral and ipsilateral hemispheres (p < 0.05).

| REAGENT or RESOURCE |  | SOURCE | | IDENTIFIER |
| --- | --- | --- | --- | --- |
| Deposited Data | |  | |  |
| Raw data | | This paper | | https://osf.io/z9rym/ |
| Software and Algorithms | | | | |
| MATLAB | | MathWorks, Natick, MA |  | https://www.mathworks.com/products/MATLAB.html, RRID: SCR_001622 |
| EEGLAB Toolbox | | Delorme & Makeig, 2004 | | https://sccn.ucsd.edu/eeglab/index.php, RRID: SCR_007292 |
| ERPLAB Toolbox | | Lopez-Calderon & Luck, 2014 | | https://github.com/lucklab/erplab/ |
| Brainstorm | | Tadel et al., 2011 | | https://neuroimage.usc.edu/brainstorm/ |
| Custom MATLAB code | | This paper | | https://github/Chenguang918/ |

### Table1
